# KDM5A/B promotes HIV-1 latency and KDM5 inhibitors promote HIV-1 lytic reactivation

**DOI:** 10.1101/2022.11.17.516956

**Authors:** Tai-Wei Li, Dawei Zhou, Zhenyu Wu, Guillaume N. Fiches, Xu Wang, Youngmin Park, Wei Jiang, Wen-Zhe Ho, Andrew D. Badley, Netty G. Santoso, Jun Qi, Jian Zhu

## Abstract

Combinational antiretroviral therapy (cART) effectively suppresses HIV-1 infection, replication, and pathogenesis in HIV-1 patients. However, the patient’s HIV-1 reservoir still cannot be eliminated by current cART or other therapies. One putative HIV-1 eradication strategy is “shock and kill”, which reactivates HIV-1 in latently-infected cells and induces their cytopathic effect or immune clearance to decrease the patients’ reservoir size. KDM5A and KDM5B act as the HIV-1 latency-promoting genes, decreasing the HIV-1 viral gene transcription and reactivation in infected cells. Depletion of KDM5 A/B by siRNA knockdown (KD) increases H3K4 trimethylation (H3K4me3) in HIV-1 Tat-mediated transactivation. We also found that the KDM5-specific inhibitor JQKD82 can increase H3K4me3 at the HIV-1 LTR region during HIV-1 reactivation and induce cytopathic effects. We applied the JQKD82 in combination with the non-canonical NF-κB activator AZD5582, which synergistically induced HIV-1 reactivation and cell apoptosis in HIV-1 infected cells. These results suggested that the KDM5 inhibition can be a putative HIV-1 latency-reversing strategy for the HIV-1 “shock and kill” eradication therapy.

## Introduction

Globally, over 36 million individuals are infected with human immunodeficiency virus type 1 (HIV-1), and currently, no approved medicine or therapy can eradicate latent HIV-1 proviruses in infected patients. HIV infection dramatically affects patients’ immune systems and causes acquired immunodeficiency syndrome (AIDS). Although combination antiretroviral therapy (cART) can suppress viral load below the detection limit in patients’ blood and halt disease progression, HIV-1 can still reactivate from latently infected cells in lymphoid tissue that maintain persistent infection in cART-treated patients. The promising shock-and-kill strategy of HIV-1 eradication therapy involves combining latency-reversing agents (LRAs) and cART to reactivate the latent HIV-1 in the infected reservoir cells without inducing global T cell activation. During HIV-1 reactivation, the infected cells would be eliminated by the viral cytopathic effect and the cellular immune response from the cytolytic T lymphocytes (CTL) or natural killer (NK) cells [1-4]. Unfortunately, most of the currently used LRAs, such as the histone deacetylase (HDAC) inhibitor vorinostat, can induce viral reactivation (as the shock step) but cannot decrease the HIV-1 latent reservoir size (as the kill step) in the clinical trial setting [2, 5, 6]. The possible reasons for this include that the HIV-1 reservoir cells, such as CD4^+^ T cells or macrophages, are resistant to the killing from CTLs [7-9] or adapt to avoid being killed by NK cells [10, 11]. Furthermore, the latently infected T cells can evade CTL/NK cells in the germinal follicles of lymphoid tissue [12-15]. For the future shock-and-kill strategy, we will need to focus on the kill strategies to facilitate the cell death of reservoir cells. Additionally, the putative strategy should activate the CTL/NK immunosurveillance or induce correct cytokine/chemokine to increase the killing of HIV-1 infected cells to eliminate the reservoir in the lymphoid tissue [9, 16].

Our previous research involved the CRISPR-knockout screening on latently infected cell lines to investigate novel genes that control HIV-1 latency and reactivation [17]. We found that potential latency-promoting genes (LPG) mainly participate in epigenetic regulation and chromatin modification/organization. Notably, the knockdown of certain histone demethylases targeting H3K4 or H3K36 methylation can induce HIV-1 reactivation. Also, the pan-Jumonji histone demethylase inhibitor JIB-04 [18-20] could induce HIV-1 reactivation in the infected T cells or monocytes. These results suggested that H3K4 or H3K36 methylation can be critical to HIV-1 reactivation and viral gene expressions. We also found that MINA53, which acts as the H3K36 demethylase, can promote HIV-1 latency due to decreased H3K36 trimethylation and inhibit transcription elongation during HIV-1 reactivation [21]. This research focused on other LPG candidates, H3K4 demethylases regulating HIV-1 latency/activation. H3K4 trimethylation (H3K4me3) is located mainly at the transcription start site (TSS) to associate with TAF3 in the TFIID complex [22-25] to facilitate transcription initiation. Previous studies showed that the HIV-1 long terminal repeat (LTR) promoter enriched H3K4me3 during HIV-1 Tat-mediated transactivation for viral gene transcription [26-28]. However, the detailed regulatory mechanism of H3K4me3 in HIV-1 LTR for viral gene expression remains unclear. We previously identified one H3K4me3 demethylase, lysine demethylase 5A (KDM5A), as a putative HIV-1 LPG that may decrease the H3K4me3 level of HIV-1 LTR to suppress HIV-1 reactivation. We hypothesize that if we can deplete KDM5A function through gene knockdown or pharmacology inhibition in HIV-1 latent infected cells, we can trigger HIV-1 reactivation in infected cells, which can be killed by the induced cytopathic effect or the CTL/NK cell immunosurveillance in the patient’s tissue.

KDM5A is one of the KDM5 demethylases (KDM5 A, B, C, and D), which catalyzes demethylation from H3K4me3 to H3K4me2 and then to the final product H3K4me1 [29-32]. Previously reported showed that KDM5A can induce PD-L1 expression [33, 34] that may suppress the immune response for immune escape during HIV-1 infection [35-39]. The other KDM5, KDM5B, has been identified to suppress immune sensing from the STING-GAS signaling [40] and RIG-I signaling [41] to decrease innate immunity and antiviral responses. Furthermore, KDM5B can prevent macrophages from releasing inflammatory cytokines like IL12 or TNF-α during *L. donovani* infection [42]. We hypothesize that KDM5A or KDM5B depletion in HIV-1 latently infected cells induces HIV-1 reactivation and stimulates innate immunity to antiviral responses. Furthermore, the H3K4m3 level of proapoptotic genes is usually low to prevent accidental cell death during the native state [23], and thus, KDM5A depletion can promote cell apoptosis [43, 44]. Therefore, we expect that inhibition of KDM5A or KDM5B can increase the antiviral responses and proapoptotic genes with HIV-1 reactivation in latently infected cells to eliminate the reservoir by HIV-1 induced cytopathic effects or antiviral-mediated programmed cell death. Above all, this study focuses on the putative KDM5 depletion treatment to achieve the shock-and-kill strategy for future putative HIV-1 eradication therapy.

## Results

### Knockdown of KDM5A increases the HIV-1 Tat/LTR-driven transcription

To identify whether KDM5A depletion increases HIV-1 LTR transcription, we performed KDM5A knockdown (KD) by siRNA in TZM-bl cells, which contain the HIV-1 LTR-driven luciferase reporter [45-47]. First, we used immunoblotting (IB) to identify decreased KDM5A expression with two specific siRNAs for knockdown (**Fig 1A**). Both KDM5A siRNAs decreased KDM5A expression in transfected TZM-bl while increasing the H3K4me3 level and IRF3 phosphorylation. Previous studies also suggested that depletion of the KDM5 family (specifically, KDM5B) can increase the IRF3 phosphorylation and transactivation to induce Type I interferon and ISG expression for antiviral responses [40, 41, 48], which can promote cell apoptosis of HIV-1 infected cells [49-52]. Also, the phosphorylated IRF3 can associate caspase-8 with Bax, promoting the proapoptotic state for cell death [53]. We used the luciferase reporter assay to identify whether the KDM5A KD affects HIV-1 LTR-driven transcription with or without HIV-1 Tat mediation (**Fig 1B**). We found that KD of siKDM5A#1 increases HIV-1 LTR transcription in the native state of TZM-bl cells. Furthermore, the KD from two different siKDM5A increased HIV-1 Tat-mediated LTR transcription in transfected TZM-bl cells. We also found that KDM5B KD increases the H3K4me3 level, phosphorylation of IRF3 (**Fig 1C**), and HIV-1 Tat-mediated LTR transcription (**Fig 1D**), consistent with the results of KDM5A KD in transfected TZM-bl cells. These results suggested that decreasing the KDM5 family, such as KDM5A or KDM5B, can increase HIV-1 reactivation and induce the IRF3-mediated proapoptotic state.

**Fig 1.**
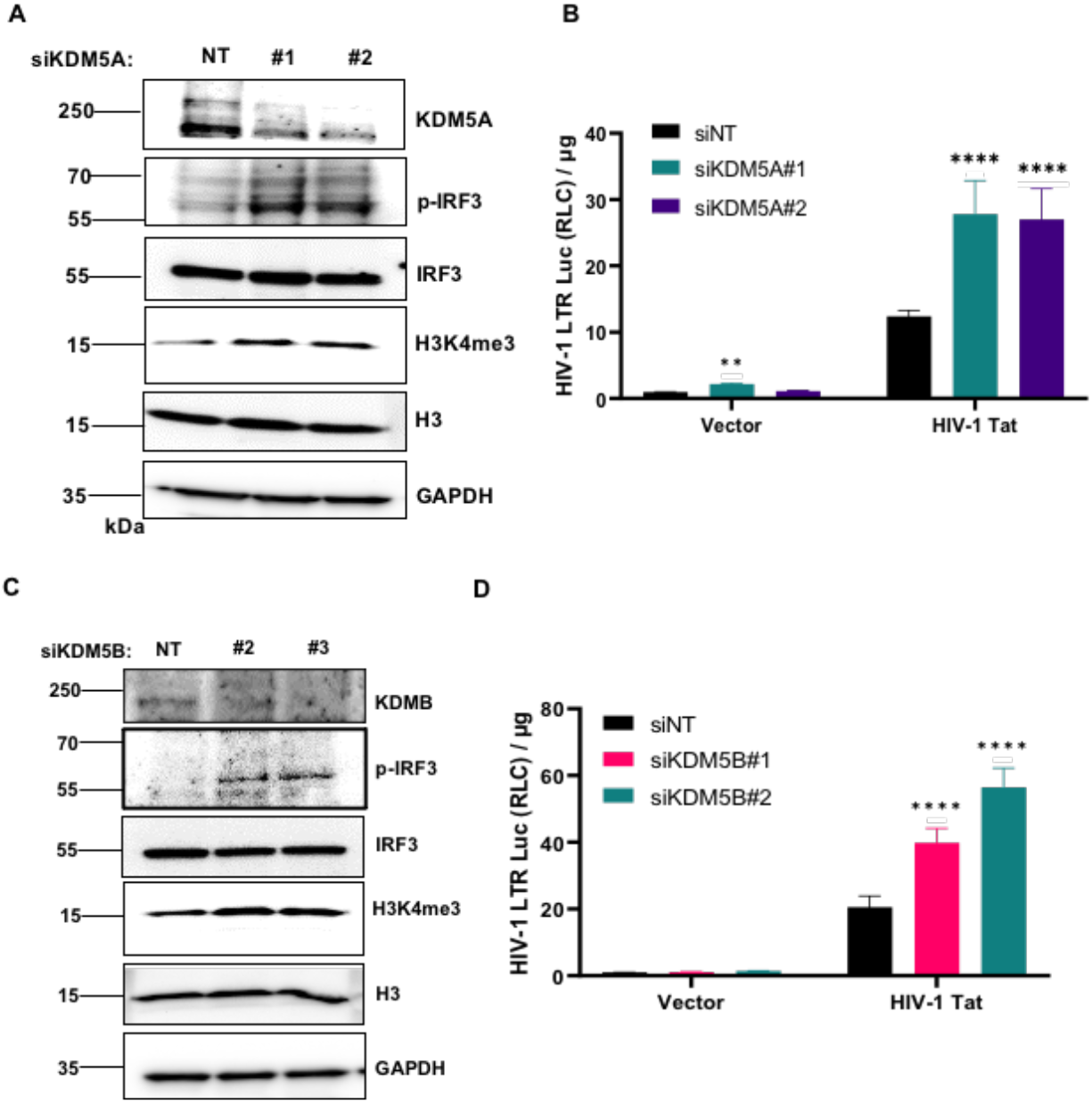
siRNA Knockdown of KDM5A/B increases the HIV-1 Tat/LTR-driven transcription. TZM-bl cells were received from reverse transfection by KDM5A siRNA # 1-2 or KDM5B siRNA #1-2 for 48h, and cells were trypsinized and reverse transfected by pQC empty vector or pQC-HIV-1 Tat for 48h. (**A**) KDM5A siRNA KD cells were harvested and lysed for IB of anti-KDM5A, p-IRF3/IRF3, GAPDH, and H3K4me3/H3. (**B**) KDM5A siRNA KD Cells with or without HIV-1 Tat overexpression were harvested for luciferase reporter assay. (**C**) KDM5B siRNA KD cells were harvested and lysed for IB of anti-KDM5B, p-IRF3/IRF, GAPDH, and H3K4me3/H3. (**D**) KDM5B siRNA KD cells with or without HIV-1 Tat overexpression were harvested for luciferase reporter assay. The readouts of RLU/total protein input (μg) were normalized with the siNT/pQC-empty vector-transfected TZM-bl control group. Results were calculated from at least 3 independent experiments and presented as mean +/-standard error of the mean (SEM). (** p <0.01; **** p<0.0001 by two-way ANOVA and Tukey’s multiple comparison test compared to the same treated siNT control).

### KDM5 inhibitor JQKD82 increases HIV-1 reactivation and cell death in latently infected cells

Since we showed that KDM5A and KDM5B KD both can promote HIV-1 LTR/Tat-mediated transcription in TZM-bl cells, we went further to investigate whether the inhibition of all KDM5 family members (such as KDM5 A, B, C, and D) [30] can promote HIV-1 reactivation in latently infected cells. We used the KDM5 inhibitor JQKD82 [44] to treat the CA5 cell line, the T lymphocyte cell line integrated with the full-length HIV-1 proviral genome and the HIV-1 LTR-driven GFP reporter [54, 55]. JQKD82 is the prodrug of KDM5-C49 [56] with the ester metabolized group to inhibit KDM5 demethylase activity with high cellular permeability. We treated the CA5 cells with a low concentration (10 or 25 μM) for 5 days to change the landscape of histone methylation and epigenetic regulation for HIV-1 reactivation. The JQKD82-treated CA5 cells showed a significant increase in the GFP expression from HIV-1 reactivation compared to the DMSO solvent control group (**Fig 2A-B**). These results showed that inhibiting KDM5 demethylase activity can directly induce HIV-1 reactivation in latently infected cells. To identify whether KDM5 inhibitor JQKD82 can increase the cell death of HIV-1 infected cells, we treated CA5 cells with JQKD82 for 5 days and stained them with LIVE/DEAD far-red dye to quantify the cell death from the drug effect or HIV-1-induced cytopathic effect [57]. CA5 cells treated with 25 μM JQKD82 had higher rates of cell death than untreated CA5 cells or Jurkat parental cells with the same treatment (**Fig 2C**). The results suggested that the inhibition of KDM5 can decrease cell survival and increase cytopathic effects in HIV-1 latently infected cells.

**Fig 2.**
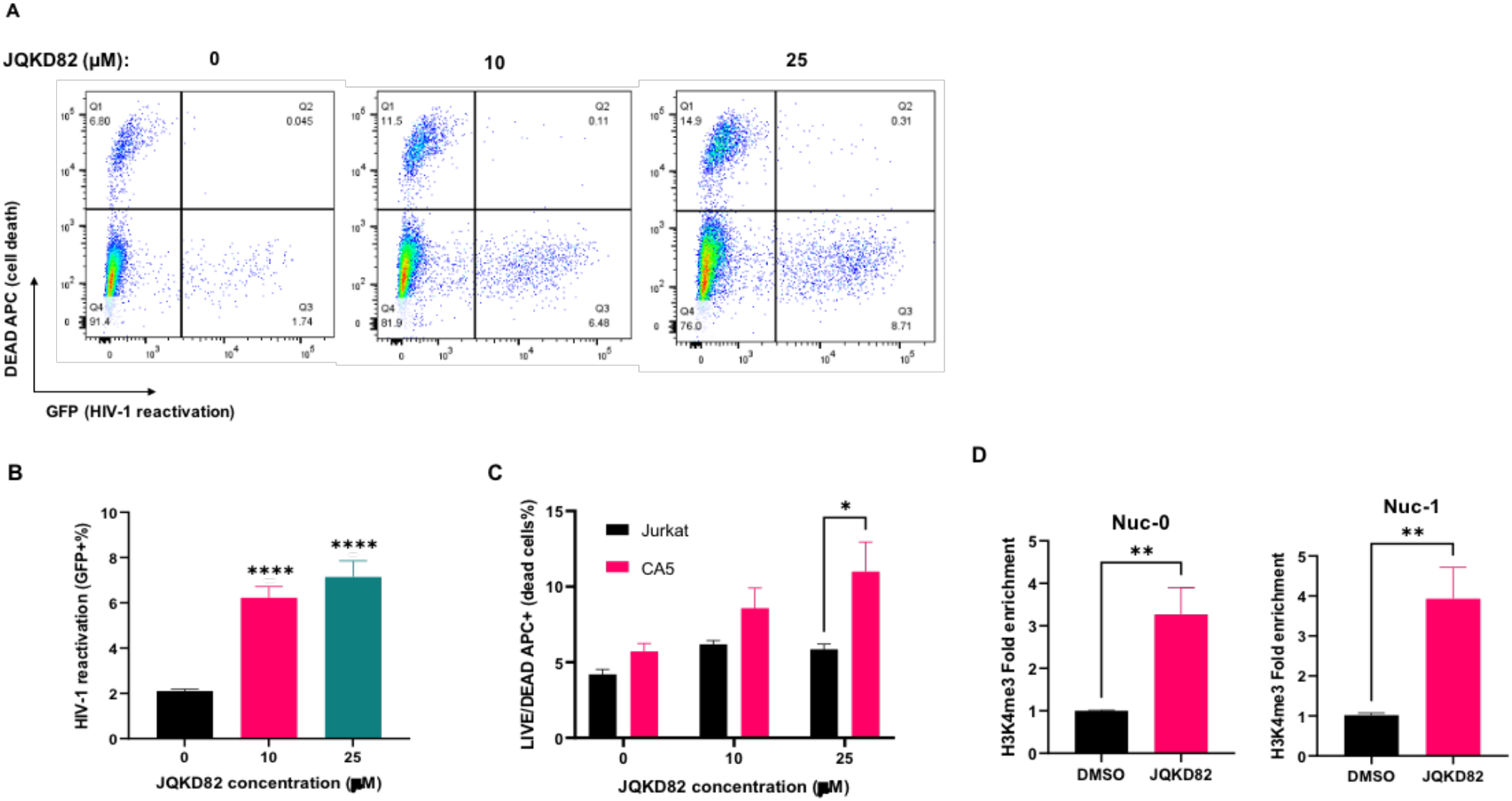
KDM5 inhibitor JQKD82 induces the HIV-1 reactivation, cell death, and H3K4me3 at HIV-1 LTR in the latent infected CA5 cells. (**A**) CA5 cells were treated with 0, 10, or 25 μM JQKD82 for 5 days. Cells were harvested for LIVE/DEAD staining and analyzed with FASC to identify the expression of HIV-1 LTR-driven GFP (**B**) and LIVE/DEAD-APC (**C**). CA5 cells were treated with DMSO or 25 μM JQKD82 for 5 days and then harvested to perform the ChIP qPCR assay of H3K4me3 (**D**) focusing on HIV-1 LTR Nuc-0 and Nuc-1 sites. Results were calculated from at least 3 independent experiments and presented as mean +/-standard error of the mean (SEM). (*p <0.05; ** p <0.01; **** p <0.0001 by one-way/two-way ANOVA and Tukey’s multiple comparison test compared to untreated (**B, D**) or parental cells control group (**C**)).

We used the chromatin immunoprecipitation (ChIP) qPCR assay to identify whether JQKD82 can increase the H3K4me3 level in the HIV-1 LTR to increase HIV-1 gene transcription for reactivation [26]. We treated CA5 cells for 5 days with DMSO or 25 μM JQKD82 and then harvested HIV-1 LTR-associated nucleosomes from cell lysates by the pull-down of anti-KDM5A or anti-H3K4me3 antibodies. We extracted the nucleosome-associated DNA and performed qPCR with the specific primers targeting the nucleosome binding sites in HIV-1 LTR (Nuc-0 and Nuc-1) [17, 58, 59]. The results showed that JQKD82 increases the H3K4me3 level in Nuc-0 and Nuc-1 (**Fig 2D**) at the HIV-1 LTR in CA5 cells compared to the untreated control group. The increase of H3K4me3 in HIV-1 LTR, especially at the Nuc-1 site where the TSS of HIV-1 viral genes [23, 24], can promote the transcription initiation to increase HIV-1 viral gene expressions and reactivation in the latently infected cells.

Also, we treated CA5 cells with a high dose of JQKD82 (50 or 100 μM) in a short period (48h), which also promoted a significant increase in HIV-1 reactivation (**Fig S1 A-B**). The high dose of JQKD82 induced significantly higher cell death of CA5 than untreated CA5 cells and Jurkat parental cells under the same treatment (**Fig S1C**). These results suggested that a high dose of JQKD82 treatment can induce HIV-1 reactivation and cell death of latently infected cells in a short period.

### JQKD82 in combination with AZD5582, increases HIV-1 reactivation and cell death in latently infected cells

Previous studies indicated that the non-canonical NF-κB activator AZD5582 (AZD) increases HIV-1 reactivation in ex vivo patient samples, humanized mouse models, and SIV-infected macaque models [60]. However, AZD5582 could not decrease the HIV-1 reservoir size in SIV-infected macaques. We hypothesized that combining JQKD82 and AZD5582 boosts HIV-1 reaction and induces cell death for the shock-and-kill strategy for HIV-1 eradication therapy. We treated CA5 cells with JQKD82 for 3 days and then refreshed the treatment with or without 0.2 μM AZD5582 for 2 days. Treated cells were harvested, and LIVE/DEAD staining was performed; using FASC, the cells were then analyzed for HIV-1 reactivation from GFP expression and cell death from APC dye signals (**Fig 3A**). AZD5582 induced ∼20% GFP expression from the treated CA5 cells, and JQKD82 further strengthened the AZD5582-mediated HIV-1 reactivation in the treated CA5 cells (**Fig 3B**). Furthermore, JQKD82 significantly increased the cell death of AZD5582-reactivated CA5 cells compared to the Jurkat cells with the same combination treatment (**Fig 3C**). We used an IB assay to identify the protein markers for cell responses from the JQKD82/AZD5582-treated Jurkat and CA5 cells (**Fig 3D**). We found that JQKD82 increases the total H3K4me3 level in both Jurkat and CA5 cells, and JQKD82 can directly increase the HIV-1 p55/p24 expression in the treated CA5 cells. With the JQKD82/AZD5582 combination treatment, JQKD82 increased the HIV-1 p55/p24 expression and the PARP cleavage (as the apoptosis marker [61, 62]) in the AZD5582-reactivated CA5 cells. We also applied a high dose of JQKD82 cotreatment with AZD5582 to Jurkat and CA5 cells for 48 hours (**Fig S2A**). The JQKD82/AZD5582 cotreatment increased the HIV-1 reactivation significantly more than a single JQKD82 or AZD5582 treatment (**Fig S2B**). The JQKD82/AZD5582 cotreatment also induced higher cell death in CA5 cells than single-drug treated CA5 cells and the JQKD82/AZD cotreat-Jurkat cells (**Fig S2C**). However, the JQKD82/AZD5582 cotreatment also caused high cytotoxicity in Jurkat parental cells (∼20% cell death).

**Fig 3.**
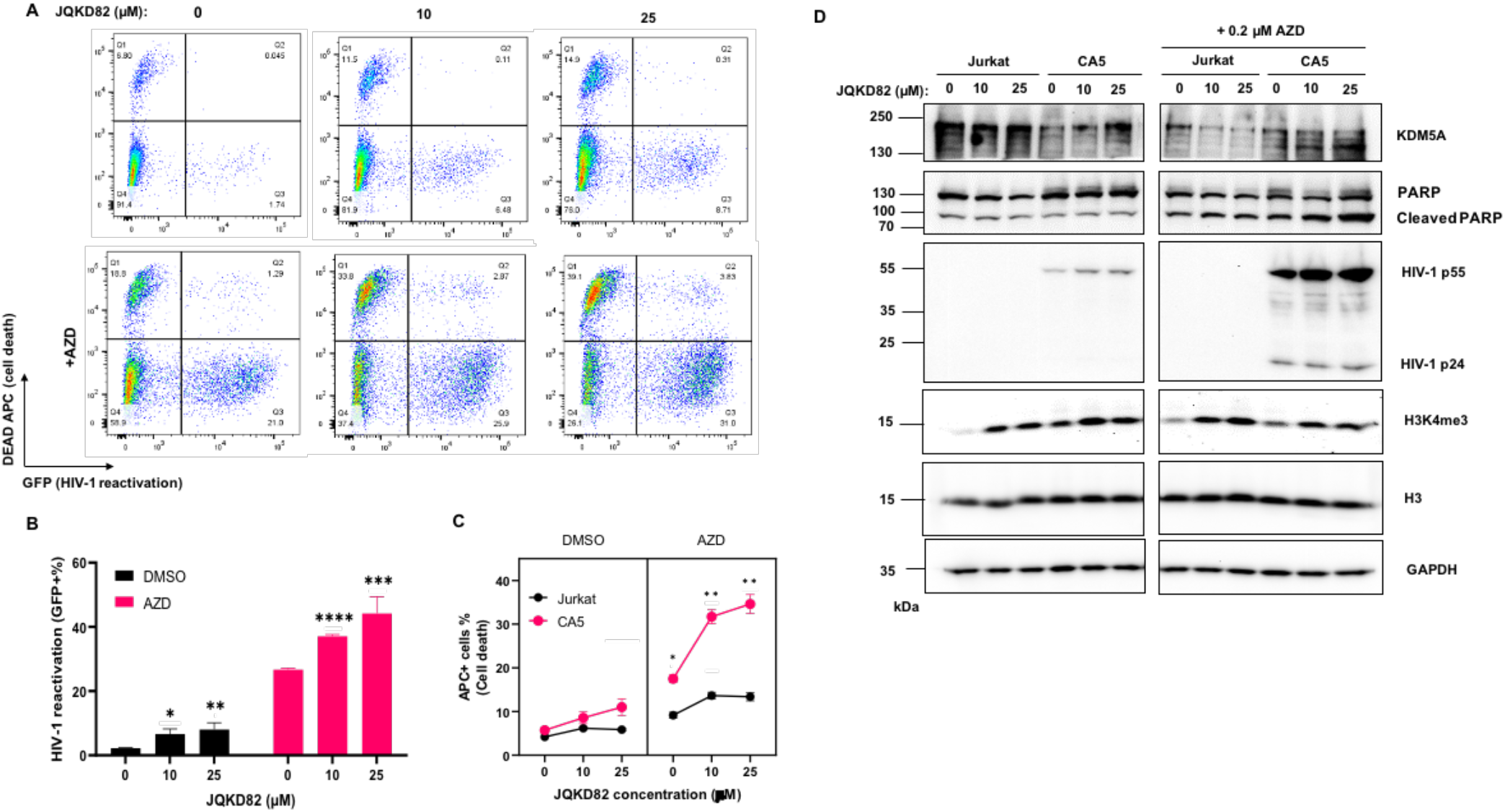
JQKD82/AZD5582 combination treatment synergistically increases the HIV-1 reactivation and cell death in CA5 cells. (**A**) CA5 cells were treated with 0, 10, or 25 μM JQKD82 for 3 days and refreshed the JQKD82-treated medium with or without 0.2 μM AZD5582 for 48h. Cells were performed the LIVE/DEAD staining and analyzed by FASC. (**B**) Treated CA5 cells were analyzed by the GFP expression from the HIV-1 reactivation. Results were calculated from 3 independent experiments and were presented as mean +/-standard error of the mean (SEM). (*p <0.05; ** p <0.01; *** p < 0.001; **** p <0.0001 by 2-way ANOVA and Tukey’s multiple comparison test compared to 0 μM JQKD82-treated control). (**C**) Treated CA5 and parental Jurkat cells were analyzed for the LIVE/DEAD APC expression. Results were calculated from 3 independent experiments and were presented as mean +/-standard error of the mean (SEM). (*p <0.05; ** p <0.01 by 3-way ANOVA and Tukey’s multiple comparison test compared to Jurkat cells under the same treatment.). (**D**) Jurkat and CA5 cells were treated with JQKD82/AZD and harvested for immunoblotting.

We treated CA5 cells with another prodrug of KDM5-C49, KDM5-C70 [56], cotreated with AZD5582 for 48h. The results suggested that the high concentration of KDM5-C70 can induce HIV-1 reactivation in CA5 cells (**Fig S3A-B**). Also, KDM5-C70 synergistically increased AZD5582-induced HIV-1 reactivation in CA5 cells. However, KDM5-C70 did not significantly increase the cell-killing effect in CA5 cells with or without AZD5582 treatment (**Fig S3C**) compared to JQKD82, which has better cell permeability than KDM5-C70 [44].

We performed the DHIV-1 infected primary Tcm model with JQKD82/AZD5582 combination treatment. We identified that the JQK82 and JQKD82/AZD5582 treatment could increase the cellular HIV-1 Gag mRNA level in the primary HIV-1 infected Tcm cells compared to the DMSO control group (**Fig 4A**). We also treated the s peripheral blood mononuclear cells (PBMCs with CD8^+^ T cell depletion) from HIV-1 aviremic patients with the JQKD82/AZD5582 combination treatment and harvested the cultured supernatant to detect HIV-1 viral RNA release. The results showed that the JQKD82/AZD5582 combination treatment increased HIV-1 viral RNA release from infected PBMCs in 4 donors (**Fig 4B**), suggesting that JQKD82/AZD5582 can induce HIV-1 reactivation in HIV-1 aviremic patient’ PBMCs. In conclusion, JQKD82 can strengthen AZD5582-mediated HIV-1 reactivation and cell killing in HIV-1 latent infected cells. A low dose of JQKD82, combined with AZD5582, can be a gradual HIV-1 eradication therapy for decreasing the latently infected cell population and HIV-1 reservoir.

**Fig 4.**
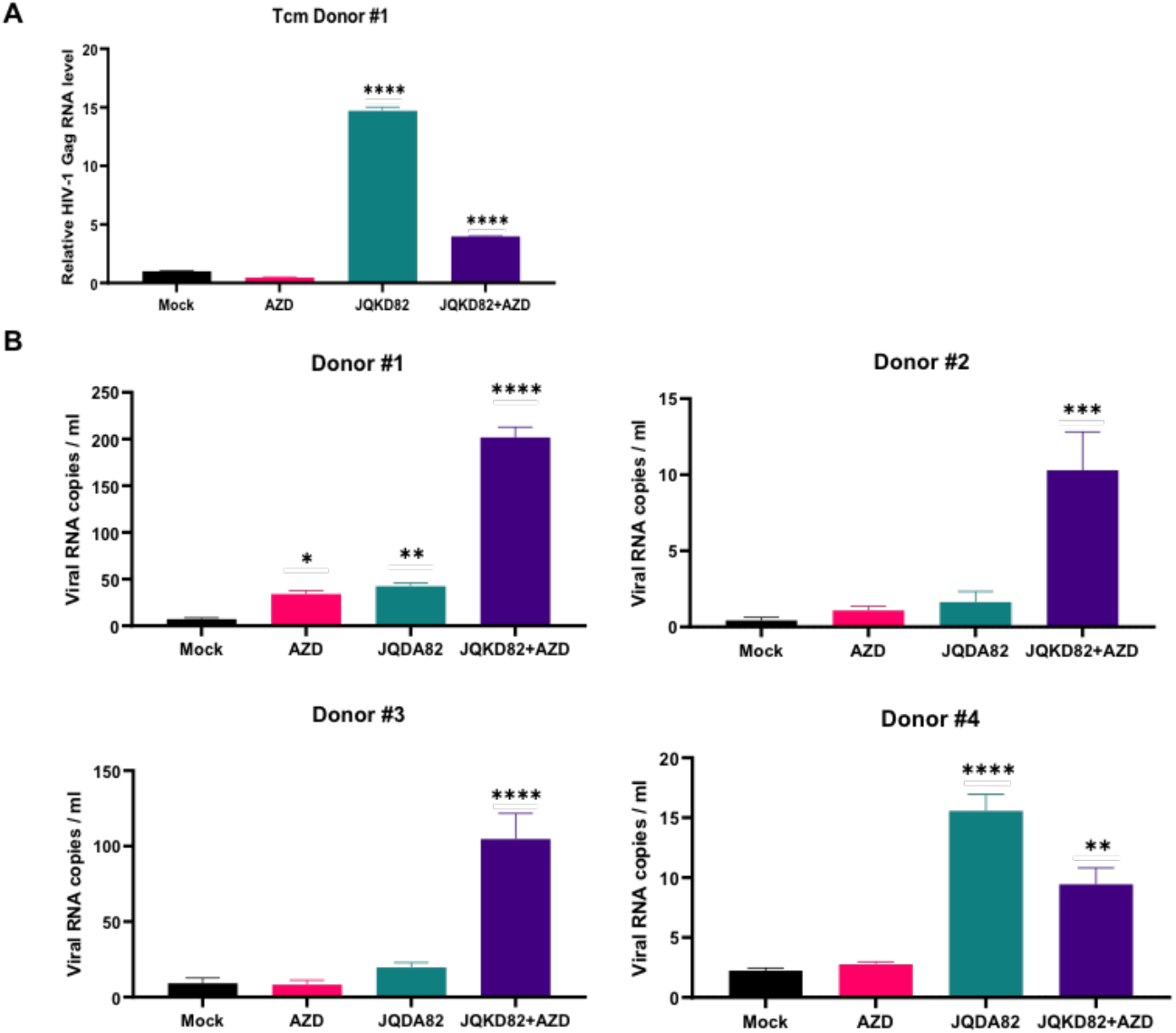
JQKD82/AZD5582 combinatory treatment induced HIV-1 reactivation in the primary T cells and HIV-1 patients’ PBMCs. (**A**) The primary Tcm latency model was established by DHIV-1 infection. The DHIV-1 infected Tcm cells were back to latency and treated with DMSO or 10 μM JQKD82 for 3 days and then refreshed the treated medium for JQKD82-treated medium for an additional 3 days with or without 0.1 μM AZD5582. Cells were harvested for RNA extraction and RT-qPCR analysis for HIV-1 Gag mRNA level. (**B**) The HIV-1 patients’ PBMC with CD8^+^ T cell-depletion for Donor #1-4 and treated with DMSO or 10 μM JQKD82 for 3 days. Then these cells were refreshed with the JQKD82-treated medium for an additional 3 days with or without 0.1 μM AZD5582. Cultured supernatant was harvested for viral RNA extraction and ultrasensitive nested RT-qPCR analysis and normalized the viral RNA copies with HIV-1 III titration standard curve. Results were calculated from 3 technical repeats and presented as mean +/-standard error of the mean (SEM). (* p<0.05, ** p <0.01; **** p <0.0001 by one-way ANOVA and Tukey’s multiple comparison test compared higher to mock control group).

### Inhibition of KDM5 A/B increases HIV-1 LTR/Tat-mediated transcription in other types of cells

We identified that the inhibition of KDM5 increases HIV-1 reactivation in latently infected CD4^+^ T cells, and we investigated whether the KDM5s regulate the HIV-1 reactivation in the monocyte/macrophage reservoirs. We used HC69 microglia cells containing HIV-1 LTR/Tat-driven GFP reporter, mimicking the cell behavior of the HIV-1 latency reservoir in the central nerve system (CNS) [63-65]. We treated HC69 cells with a KDM5 inhibitor, JQKD82, for 5 days and found that JQKD82 significantly increased HIV-1 LTR/Tat-driven GFP expression (**Fig 5A-B**), but JQKD82 did not significantly increase the cell death of HC69 cells (**Fig 5C**). We also used siRNA KD of KDM5A or KDM5B to identify whether H3K4me3 demethylase promotes HIV-1 latency in infected microglia. The KD of KDM5A or KDM5B significantly increased HIV-1 LTR/Tat-driven GFP in transfected HC69 cells (**Fig 5 D-E**) but did not cause significant cell death in microglia cells (**Fig 5F**). These results suggested that inhibition of KDM5A or KDM5B can increase HIV-1 reactivation in the monocyte/macrophage reservoir and need to be combined with other reagents or interventions to eradicate HIV-1 infected microglia cells in CNS.

**Fig 5.**
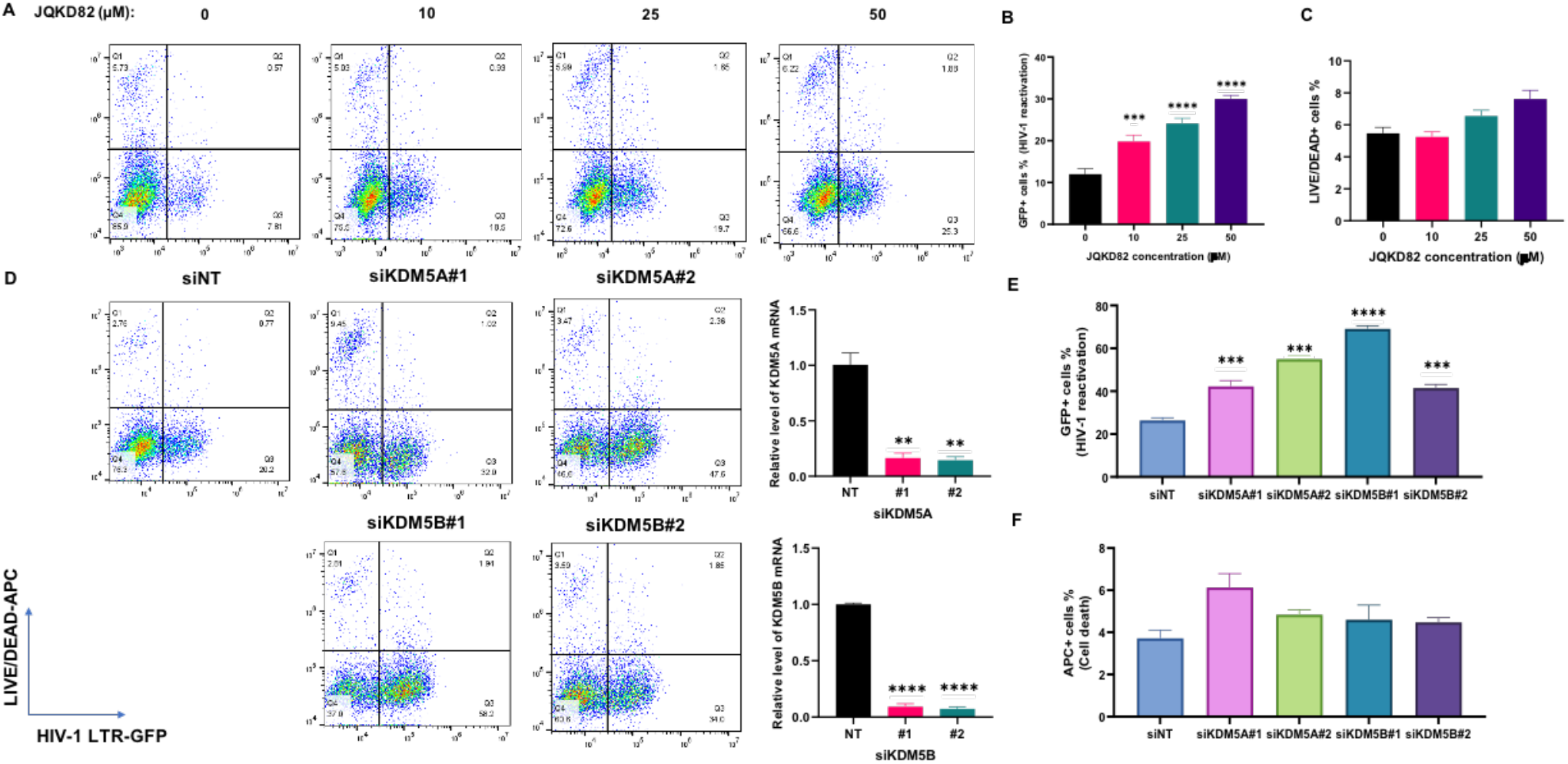
KDM5 inhibitor JQKD82 or siRNA knockdown of KDM5A/B induced the HIV-1 latency reactivation in HC69 microglia cells. (**A**) HC69 microglial cells were treated with 0, 10, 25, or 50 μM JQKD82 for 5 days. Cells were harvested for LIVE/DEAD staining and analyzed with FASC to identify the expression of HIV-1 LTR-driven GFP (**B**) and LIVE/DEAD-APC (**C**). HC69 microglia cells were received from reverse transfection by KDM5A siRNA # 1-2, or KDM5B siRNA #1-2 for 72hr. The siRNA-transfected cells were harvested for RNA extraction and RT-qPCR to identify the siRNA KD efficiency. Cells were harvested for LIVE/DEAD staining and analyzed with FASC to identify the expression of HIV-1 LTR-driven GFP (**E**) and cell death (**F**). Results were calculated from 3 independent experiments and presented as mean +/-standard error of the mean (SEM). (*** p <0.001; **** p <0.0001 by one-way ANOVA and Tukey’s multiple comparison test compared to the untreated or SiNT control group).

We also gave JQKD82/AZD5582 combination treatment to U1/HIV cells, the monocytic cell line latently infected by HIV-1 [66-68], for 5 days (**Fig S4A**). The results showed that JQKD82/AZD5582 combination treatment significantly increases HIV-1 Gag protein expression (**Fig S4B)** and mRNA level (**Fig S4C**). These results suggested that JQKD82/AZD5582 combination treatment induces HIV-1 reactivation in latently infected monocytes.

## Discussion

The results of this study suggested that the depletion of KDM5A or KDM5B can raise HIV-1 Tat/LTR mediated transcription to promote HIV-1 reactivation (**Fig 1**). We hypothesized that KDM5s (Specially KDM5A) can recognize the H3K4me3 at HIV-1 LTR and catalyze the H3K4 demethylation to increase the threshold for reactivation and promote latency in HIV-1 infected cells. After KDM5 depletion increases in the H3K4me3 level at HIV-1 LTR sites, H3K4me3 facilitates the transcription initiation and HIV-1 Tat/TAR-mediated reactivation. We also found that the KD of KDM5A/B is insufficient to turn on HIV-1 LTR transcription in TZM-bl cells directly. However, KDM5-inhibitor treatment may increase other viral protein expressions, such as HIV-1 Tat, as positive feedback to facilitate the HIV-1 reactivation in the HIV-1 latent CA5 cells (**Fig 2**). In conclusion, the axis of KDM5/H3K4me3/Tat interaction to regulate the transactivation of HIV-1 LTR promoter can be critical for the epigenetic controls of HIV-1 reactivation and latency in latently infected cells.

In previous studies about the epigenetic control of HIV-1 reactivation, HDAC inhibitors such as valproic acid [69, 70], panobinostat, or SAHA (Vorinostat) [71-73] were used, which could induce HIV-1 reactivation in the latently infected cell line, primary infected cell model, or even in ex vivo aviremic ART-treated patients’ infected cells. However, these HDAC inhibitors could not pass human clinical as they were being insufficient to progressively reactivate HIV-1 latent infected cells in the lymphoid tissue over a long period [5, 74], failing to terminate HIV-1 infection in resting T cells during HIV-1 reservoir expansion [75, 76], or limiting the cell killing effect [6, 77-80] to decrease the reservoir in the patient’s body [81]. The future shock-and-kill therapy would need to pursue or combine the different putative drugs to enhance reactivation and promote the killing of HIV-1 reservoirs. Previous studies suggested that KDM5A and KDM5B can associate with HDAC-complex to perform epigenetic silencing cooperatively [82-85]. KDM5 inhibitors combined with HDAC inhibitors could benefit future HIV-1 eradication by improving the reactivation and cell-killing effects. Also, the depletion of KDM5B can potentially increase antiviral immunity and promote IRF3 signaling (**Fig 1C**), which can increase the IRF3-mediated or antiviral-induced cell death [53, 86-88] to decrease the HIV-1 reservoir expansion and archive the shock-and-kill therapeutic process.

JQKD82 is the prodrug with the ester-modified group of the functional metabolite KDM5-C49, which can block the α-ketoglutarate catalytic site in the KDM5 Jumonji-C domain [44, 56]. In a previous study, JQKD82 showed better cell permeability and cellular accumulation than other C49-derivatives or prodrugs and significantly increased the global H3K4me3 level in treated cells. In RNA-Seq analysis, the gene expressions after JQKD82-treatment increased in Type I interferon and inflammatory responses [44]. Like other KDM5 inhibitors and depletions, JQKD82 can cause the shut-down cell cycle and promote the proapoptotic state and innate immunity responses [43, 44, 89, 90]. We applied the low doses of JQKD82 treatment (below 25 μM) for an extended period (at least 5 days) to HIV-1 latent cells and induced HIV-1 reactivation and their cell death (**Fig 2**). We also applied the high doses of JQKD82 treatment (beyond 50 μM) to latently infected cells and uninfected parental cells for a short time (48h). This treatment induced HIV-1 reactivation but also caused high cell-killing effects in uninfected parental cells (**Fig S1C**). These results suggested that intense JQKD82 can cause high cytotoxicity in uninfected cells.

In this study, we used the KDM5 inhibitor, JQKD82, in combination with the non-canonical NF-κB activator, AZD5582, to synergistically increase HIV-1 reactivation and cell death in latently infected cells. AZD5582 is a SMAC-mimetic analog that causes inhibitor of apoptosis proteins (IAPs) self-ubiquitination and degradation [60, 91] to increase p52-RelB nuclear translocation and turn on downstream gene expressions [92]. AZD5582 and other SMAC-mimetic IAP inhibitors can also increase the cellular proapoptotic state for induced HIV-1 cytopathic killing [93-96]. AZD5582 could reactivate HIV-1/SIV in patient cell samples and animal models [60]. However, AZD5582 alone could not decrease the reservoir size in SIV-infected macaques and increase antiviral immunity responses. Our results showed that the combination of JQKD82 and AZD5582 treatment boosted the HIV-1 reactivation and induced apoptosis (**Fig 3**) in the latently infected cells and did not cause severe cytotoxicity in the uninfected parental cells. We also found that another KDM5-C49 prodrug, KDM5-C70, can synergetically increase AZD5582-induced HIV-1 reactivation, but KDM5-C70 cannot increase the cell-killing effect as JQKD82 treatment (**Fig S3**). These results suggested that JQKD82 has a better cell permeability and KDM5-inhibitory effect [44], promoting a higher proapoptotic state in HIV-1 latently infected CD4^+^ T cells and monocytes. We also performed JQKD82/AZD5582 combination treatment to the HIV-1 patient’s PBMCs (with the depletion of CD8^+^ T cells) and found that JQKD82/AZD5582 can induce HIV-1 reactivation and increase the release of HIV-1 viral RNA from 4 donors’ infected PBMC (**Fig 4B**). In the future, we will test more PBMC samples from different HIV-1 patients to confirm these preclinical results. Also, we will quantify the change in reservoir sizes of JQKD82/AZD5582-treated HIV-1 infected PBMCs by the quantitative viral outgrowth assay (QVOA) to validate whether the JQKD82/AZD5582 combination treatment can be the candidate for shock-and-kill therapy.

HIV-1 infected microglia cells are the latency reservoir and cause the abnormal inflammatory state for the pathogenesis of HIV-1-associated neurocognitive disorder (HAND), even under the cART control [97]. In order to understand the HIV-1 reactivation mechanism of microglia and develop a therapy targeting HIV-1 reservoirs in the CNS, we used the immortalizing human primary microglia with HIV-1 LTR/Tat-mediated GFP reporter, HC69 cell line [63-65] to investigate whether the KDM5 depletion can increase HIV-1 reactivation and cytopathic effect. The JQKD82 treatment and KDM5 A/B siRNA KD can increase the HIV-1 LTR/Tat transactivation in HC69 microglia (**Fig 3.5**). However, the depletion of KDM5, whether by pharmaceutical inhibitors or siRNA knockdown, can not increase the cytopathic effect or cell death in HC69 microglia. HC69 cells contain the HIV-1 LTR/Tat driven GFP reporter, but HC69 cells have the deletion of HIV-1 Gag, Pol, and other accessory viral proteins, which cause single-round infection without generating severe cytopathic effects during HIV-1 reactivation. These results suggested that the KDM5 inhibitors can be a putative LRA for the HIV-1 reactivation in infected microglia, but we need to identify their cell-killing effects in the different HIV-1 infected microglia models.

In this study, we demonstrated that the depletion of KDM5 A/B by siRNA KD or enzymatic inhibitor JQKD82 increases the H3K4me3 at the HIV-1 LTR site and HIV-1 Tat-mediated viral gene transcriptions for HIV-1 reactivation. The KDM5 depletion combination with non-canonical NF-κB activator AZD5582 can significantly increase cell death in HIV-1 infected cells. The JQKD82/AZD5582 combination treatment can promote the HIV-1 reactivation and proapoptotic state to eliminate the HIV-1reservoir for putative HIV-1 cure therapy.

### Material and methods Cell culture

TZM-bl, Jurkat, and U1 cell lines were obtained from the National Institutes of Health (NIH) AIDS reagent program. The Jurkat-derived CA5 cell line latently infected with replication-competent, full-length HIV-1 genome was provided by Dr. O. Kutsch [54, 55]. TZM-bl and HEK293T (Cat. # CRL-3216, ATCC) cells were cultured in Dulbecco’s modified Eagle’s medium (DMEM, Cat # D5796, Sigma). All T cell lines were cultured in Roswell Park Memorial Institute (RPMI) 1640 medium (Cat # 11875093, Gibco). Completed cell culture medium contained 10% fetal bovine serum (FBS, Cat. # 10437028, Thermo Fisher), penicillin (100 U/ml) /streptomycin (100 μg/ml) (Cat. # MT30002CI, Corning). Primary peripheral blood mononuclear cells (PBMCs) were maintained and cultured with a completed RPMI medium with 1× minimum essential medium nonessential amino acid (Cat #11140-050, Gibco), 1× sodium pyruvate (Cat #11360-070, Gibco), and 20 mM HEPES (Cat #15630-080, Gibco). Human recombinant IL-2 (rIL-2, Roche) at 30 U/ml was supplied to primary cells every 2 days [57]. HC69 and the parental C20 microglia cells were cultured in BrainPhys medium (Cat. #05790, StemCell Technologies) containing N2 supplement (Cat. #17502048, Gibco), penicillin (100 U/ml) /streptomycin, 100 μg/mL normocin (Cat. #ant-nr-1, InvivoGen), 25 mM glutamine (Cat. #25030081, Gibco), 1% FBS and 1 μM DEXA (Cat. #D4902, Sigma) freshly added to the cell culture [63-65].

### Compounds, antibodies, and plasmid

Recombinant human TNF-α (Cat. # 554618) was purchased from BD. Biosciences. KDM5 inhibitor JQKD82 was synthesized and generously gifted by Dr. Jun Qi’s lab [44]. AZD-5582 (Cat. # CT-A5582) was purchased from Chemie Tek. KDM5-C70 (M60192-10S) was purchased from Xcess Biosciences.

Mouse anti-GAPDH antibody (Cat. # sc-32233) was purchased from Santa Cruz Biotechnology. Rabbit anti-phosphorylated IRF3 antibody (Cat. # 4947S), rabbit anti-IRF3 antibody (Cat. # 4302S), rabbit anti-H3 antibody (Cat. # 9715S), rabbit anti-PARP antibody (Cat. # 9542T), goat HRP-conjugated anti-mouse IgG antibody (Cat. # 7076S), and goat HRP-conjugated anti-rabbit IgG antibody (Cat. # 7074) were purchased from Cell Signaling Technology. Mouse anti-KDM5A antibody (Cat # 91211) and Mouse anti-H3K4me3 antibody (Cat #61379) were purchased from Active Motif. HIV-1 Gag p24 IgG1 monoclonal antibody was produced from the hybridoma cell line (NIH AIDS reagent program).

The pQC-HIV-1 Tat was constructed by subcloning C terminal Flag tag fused HIV-1 Tat into the pQCXIP retroviral empty vector (Clontech) using NotI and BamHI sites [98].

### Transient transfection

For KDM5A knockdown, 10 nM siRNA (siKDM5A; #1: siRNA ID: s11835: 5’-CCGCUAAAGUGGAAGCUAUtt-3’; #2: siRNA ID: s11836: 5’-GCGAGUUUGUUGUGACA-UUtt-3’; siKDM5B #1: siRNA ID: s21145: 5’-GGCAGUAAAGGAAAUCGAAtt ; #2: siRNA Id: s21146: 5’-GGAAGAUCUUGGACUUAUUtt-3’, Ambion by Life technologies; non-targeting control: Silencer™ Negative Control No. 4 siRNA, si N.T., Cat. # AM4641, Invitrogen) was reversely transfected in TZM-BL cells using Lipofectamine™ RNAiMAX Transfection Reagent (Cat. # 13778030, Invitrogen). Cells were kept in culture for 48h and continued the further experiment subjected to IB of KDM5A or KDM5B to evaluate the knockdown efficacy.

For HIV-1 Tat overexpression to induce the HIV-1 LTR-driven reporter assay, we performed the transient transfection of pQC-HIV-1 Tat or pQCXIP empty vector control in TZM-BL cells using Fugene6 transfection reagents (Cat. # E2691, Promega). Briefly, cells were seeded and incubated with the mixture of plasmids with Fugene 6 for 24 h, following the manufacturer’s protocol. The medium was changed and further cultured for an additional 24h for harvesting and performed following experiments.

### Luciferase reporter assays

Treated TZM-BL cells were trypsinized and harvested for luciferase activity assay (One-Glo System, Cat. # E6110, Promega) following the manufacturer’s protocol. Chemiluminescence was determined by using the was detected using Biotek Cytation5 and analyzed by GEN5 software (Biotek). In the HIV-1 LTR-driven reporter assay, the relative light unit (RLU) of luciferase luminescence was divided by the total protein input (RLU/μg) quantified by the BCA assay kit (Cat. #23225, Thermo Scientific). The readouts were normalized with the siNT/pQC-empty vector-transfected TZM-BL group.

### Protein immunoblotting

Protein immunoblotting was performed following our previously published protocols [57, 99]. Briefly, cells were harvested, washed by PBS, and pelleted. Cell pellets were lysed in RIPA buffer (Cat. #20-188, Millipore) containing protease inhibitor cocktail (Cat. # A32965, Thermo Scientific) on ice, followed by brief sonication to prepare cell lysate. The BCA assay kit (Cat. #23225, Thermo Scientific) was used to quantify the total protein amount in cell lysate, which was boiled in the SDS loading buffer with 5% β-mercaptoethanol (Cat. #60-24-2, Acros Organics). The denatured protein samples were separated by Novex™ WedgeWell™ 4-20% SDS-PAGE Tris-Glycine gel and transferred to PVDF membrane (iBlot™ 2 Transfer Stacks, Invitrogen) using iBlot 2 Dry Blotting System (Cat. # IB21001, Thermo Scientific). The membranes were blocked by 5% milk in PBST and probed by the specific primary antibodies at 4°C overnight, followed by the HRP-conjugated secondary antibodies. The membranes were developed using the Clarity Max ECL substrate (Cat. # 1705062, Bio-Rad).

### Cell viability assay

The death of HIV-1-reactivated or compound-treated cells was determined using the LIVE/DEAD Fixable Far Red Dead Cell Stain Kit (Cat. # L10120, Invitrogen), following the manufacturer’s protocol [57]. In brief, the treated cells were washed and incubated with the working dilution of LIVE/DEAD dye for 30 mins and then washed with PBS. Stained cells were fixed with 4 % paraformaldehyde (Cat. # 15714S, Electron Microscopy Sciences), and analyzed the APC signaling by the BD Accuri C6 Plus flow cytometer (BD Biosciences).

### Flow cytometry

Cells were harvested, washed twice with PBS, fixed by 4% paraformaldehyde, and then resuspended in 2% BSA with PBS for FASC analysis. The cell samples were analyzed using an Accuri C6 Plus flow cytometer (BD Biosciences) with forward versus side scatter (FSC-A versus SSC-A) gating and the corresponding optical filters for the excitation/emission of fluorescence expression. The percentage of fluorescence-positive cells was determined by using the FlowJo V10 software.

### Chromatin immunoprecipitation (CHIP) assay

ChIP assay was conducted as described previously [17, 59, 99, 100]. Cells were cross-linked by using 0.5% paraformaldehyde for 10 min, followed by treatment with 125 mM glycine to quench the reaction for 5 min. After washing with cold PBS twice, cells were lysed for 10 min on ice in CE buffer (10 mM HEPES-KOH, 60 mM KCl, 1 mM EDTA, 0.5% NP-40, 1 mM DTT, pH: 7.9 with protease inhibitor cocktail). The nuclei were pelleted by centrifugation at 700 × g for 10 min at 4°C and resuspended in SDS lysis buffer (1% SDS, 10 mM EDTA, 50 mM Tris-HCl, Ph: 8.1 with protease inhibitor cocktail). Nuclear lysates were sonicated for 2 min to fragment genomic DNA and subsequently diluted with ChIP dilution buffer (0.01% SDS, 1% Triton X-100, 1.2 mM EDTA, 16.7 mM Tris-HCl, 150 mM NaCl, pH: 8.1 with protease inhibitor cocktail). The lysates were incubated overnight at 4°C with specific antibodies or control mouse IgG (Cat. # sc-2025, Santa Cruz). Protein A/G beads (Cat. # 88803, Pierce) were pre-blocked with 0.5 mg/ml BSA and 0.125 mg/ml herring sperm DNA (Cat. 15634-017, Invitrogen) for 1 h at room temperature and then added to the lysate-antibody mixture for another incubation at 4°C for 2 h. Beads were washed with the following buffers: low salt wash buffer (0.1% SDS, 1% Triton X-100, 2 mM EDTA, 20 mM Tris-HCl, 150 mM NaCl, pH 8.1); high-salt wash buffer (0.1% SDS, 1% Triton X-100, 2 mM EDTA, 20 mM Tris-HCl, 500 mM NaCl, pH 8.1); LiCl buffer (0.25 M LiCl, 1% NP-40, 1% Na-deoxycholate, 1 mM EDTA, 10 mM Tris-HCl, pH 8.1); and TE buffer (10 mM Tris-HCl, 0.1 mM EDTA, pH 8.1), and were eluted with fresh elution buffer (1% SDS, 0.1 M NaHCO_3_) at room temperature. The eluted samples were incubated at 65°C overnight in the presence of 0.2 M NaCl to disassociate the cross-linking of protein/bound DNA. The eluted samples were then treated with proteinase K (Cat. #EO0491, Thermo Fisher Scientific) for protein digestion at 50°C for 2h, and the DNA species were precipitated by using UltraPure™ Phenol:Chloroform: Isoamyl Alcohol reagent (Cat. # 15593031, Thermo Fisher Scientific) with the manufacturer’s protocol. The DNA pellets were dried out and resuspended in water, and the pull-down DNA was quantified by qPCR.

### Quantitative reverse transcription PCR (RT-qPCR and ChIP-qPCR

RT-qPCR assays were performed following the previously published protocol [101]. Total RNAs from harvested cells were extracted using the NucleoSpin RNA extraction kit (Cat. # 740955.250, MACHEREY-NAGEL), and ∼1 μg RNA was reversely transcribed using the iScript™ cDNA Synthesis Kit (Cat. # 1708890, Bio-Rad). Real-time qPCR was conducted using the iTaq™ Universal SYBR® GreenSupermix (Cat. # 1727125, Bio-Rad). The PCR reaction was performed on a Bio-Rad C.F.X. connect qPCR machine under the following conditions: 95 °C for 10 m, 50 cycles of 95 °C for 15 s, and 60 °C for 1 m. Relative gene expression was normalized to GAPDH internal control as the 2^-ΔΔCt^ method: 2 ^(ΔCT of targeted gene - ΔCT of GAPDH)^. The qPCR primers were used in this study as below: HIV-1 Gag (F: CTGAAGCGCGCACGGCAA; R: 5’-CTGAAG CGCGCACGGCAA-3’), β actin (F: 5’-GGACCTGACTGACTACCTCAT-3’; R: 5’-GTAGCACAGCTTCTCCTTAAT-3’), KDM5A (F: 5’-CAACGGAAAGGCACTCTCTC-3’; R: 5’-CAAGGCTTCTCGAGGTTTG-3’), KDM5B (F: 5’-ATTCTGTTGGCACATTGAAGACC-3’; R: 5’-AGCATACCCTGGGACTCCATAC-3’).

ChIP-qPCR assays were performed by the elute DNA from the ChIP assay. The fold enrichment from ChIP was normalized to IgG pull-down control in the same treatment as the 2^-ΔΔCt^ method: 2 ^(ΔCT of targeted gene -ΔCT of IgG pull-down)^. The PCR primers for the bound nucleosomes of HIV-1 LTR were: Nuc-0 (F: 5’-GAAGGGCTAATTTGGTCCCA -3’; R: 5’-GATGCAGCTCTC GGGCCATG-3’), Nuc-1 (F: 5’-AGTGTGTGCCCGTCTGT-3’; R: 5’-TTGGCGTACTCACCA GTCGC-3’).

### DHIV-1 preparation

Envelope-deficient DHIV backbone plasmid was provided by Dr. Vicente Planelles [102]. VSV-G pseudo-typed DHIV viruses were prepared by transfecting HEK293T cells with DHIV vector and pMD2.G-VSV-G plasmids with Turbofect reagent (Cat. #R0531, Thermos scientific) following the manuscript protocol. The supernatant containing VSV-DHIV viral particles was harvested at 48 hours after transfection. Supernatant-containing viruses were centrifuged to remove cellular debris, filtrated with 0.45-μm membrane filters, and stored at −80°C [57]. Viral supernatant was tittered by infecting Jurkat cells with serial dilution for 48h and performing anti-HIV-1 Gag immunofluorescence staining. The percentages of HIV-1 Gag positive cells from different dilutions of supernatant were analyzed by FASC. The infectivity of the VSV-G DHIV-1 supernatant ranged from 1.7-2.9 × 10^7^ IU/ml.

### Establishment of HIV-1 latency in human primary CD4+ T cells

A primary CD4+ Tcm cell model of HIV-1 latency established by Dr. Vicente Planelles’ group was used as previously described with modifications [57, 102]. The frozen Human primary peripheral blood CD4^+^ T cells (Cat. # 200-0165, Stemcell Technologies) were stimulated on a 96-well Nunc-Immuno Maxi Sorp plate precoated with soluble anti-CD3/CD28 antibodies in RPMI complete medium containing TGF-β1 (10 ng/ml), anti-human IL-12 (2 μg/ml), and anti-human IL-4 (R&D Systems) (1 μg/ml) for 3 days. After 3-day activation, the naïve cells were considered nonpolarized, differentiated into a TCM-like phenotype, and cultured in RPMI complete medium with Human rIL-2 (30 U/ml). On day 7, the Tcm-like cells were infected with VSV-G pseudo-typed DHIV (MOI: 1.0; p24 input 100 ng/ml) through spinoculation with 1 × 10^6^ cells/ml with 8 μg/ml polybrene (Cat. # TR-1003-G, Sigma) at 1741g at 37°C for 2 hours. Infected Tcm cells were cultured for 10-day incubation to establish HIV-1 latency. KDM5 inhibitor JQKD82 (10 μM) was treated for DHIV-infected T cm for 3 day and refreshed with JQKD82-treatment with or without 0.1 μM AZD-5582. We harvested these cells for RNA extraction and RT-qPCR analysis for detecting HIV-1 Gag mRNA level (β actin mRNA as the internal control).

### Ex vivo analysis using CD8-depleted PBMCs of aviremic patients

Consented HIV-1 positive, cART-treated, aviremic patients (<20 copies per mL) were recruited through the AIDS clinic at the Mayo Clinic to donate whole blood via leukapheresis. Peripheral blood mononuclear cells (PBMCs) were then isolated and cultured in a complete medium supplemented with 30 U/mL interleukin-2 (IL2) and in the presence of 600 nM of Nevirapine (Cat. #, Sigma) for 3 days. PBMCs were subjected to CD8 ^+^ T cells depletion by negative selection using the CD8 MicroBeads (Cat. # 130-045-201, Miltenyi Biotec). CD8^+^ T cells-depleted PBMCs were treated with or without 10 μM JQKD82 for 3 days, and then the JQKD82-untreated were refreshed in the cultured medium with no-treatment (mock), 0.1 μM AZD5582 (AZD), or PMA (100 ng/ml) +ionomycin (0.5 μg/ml) or anti-CD3/CD28 Dynabeads (ratio: 1 bead to 2 cells; Cat. # 11161D, Gibco). The JQKD82-treated cells were changed medium with 10 μM JQKD82 with or without 0.1 μM AZD5582. Supernatants were collected and subjected to the extraction of HIV viral RNAs by using the QIAmp® Viral RNA Kit (Cat. #52904, Qiagen). The ultrasensitive nested qPCR assay was performed to quantify HIV viral RNA copies as previously described [100, 103, 104]. HIV-1 IIIB RNAs were extracted by the NucleoSpin RNA Virus kit (Cat. #740956.10, MACHEREY-NAGEL) and quantified with copy numbers by the Lenti-X qRT-PCR titration kit (Cat. #631235, Takara Bio). A serial dilution of HIV-1 IIIB viruses with known concentrations at a series of dilutions were used to create a standard curve for the absolute quantification of reactivated HIV-1 viruses in supernatants.

### Statistics

Statistical analysis was performed using GraphPad PRISM. Data are presented as mean ± standard error of the mean (SEM) of biological repeats from at least 3 independent experiments. * p<0.05, ** p<0.01, *** p<0.001, or **** p<0.001 indicated the significant difference analyzed by ANOVA and Tukey’s multiple comparison test.

## Acknowledgments

We thank the cooperation of and contribution of the lab of Dr. Jun Qi from Dana–Farber Cancer Institute. We thank Dr. Andrew D. Badley from Mayo Clinic for providing the HIV-1 patients’ donated PBMC. We thank Dr. Olaf Kutsch from University of Alabama at Birmingham providing the CA5 cell line. We also thank Dr. Karin Musier-Forsyth, Dr. Shan-Lu Liu, and Dr. Namal Liyanage at The Ohio State University for their advice on our studies. This study was funded by NIH research grants R01AI150448, R01DE025447, R56AI157872, and R33AI116180 to JZ; R03DE029716, R01CA260690 to NS.

---

**Fig S1.**
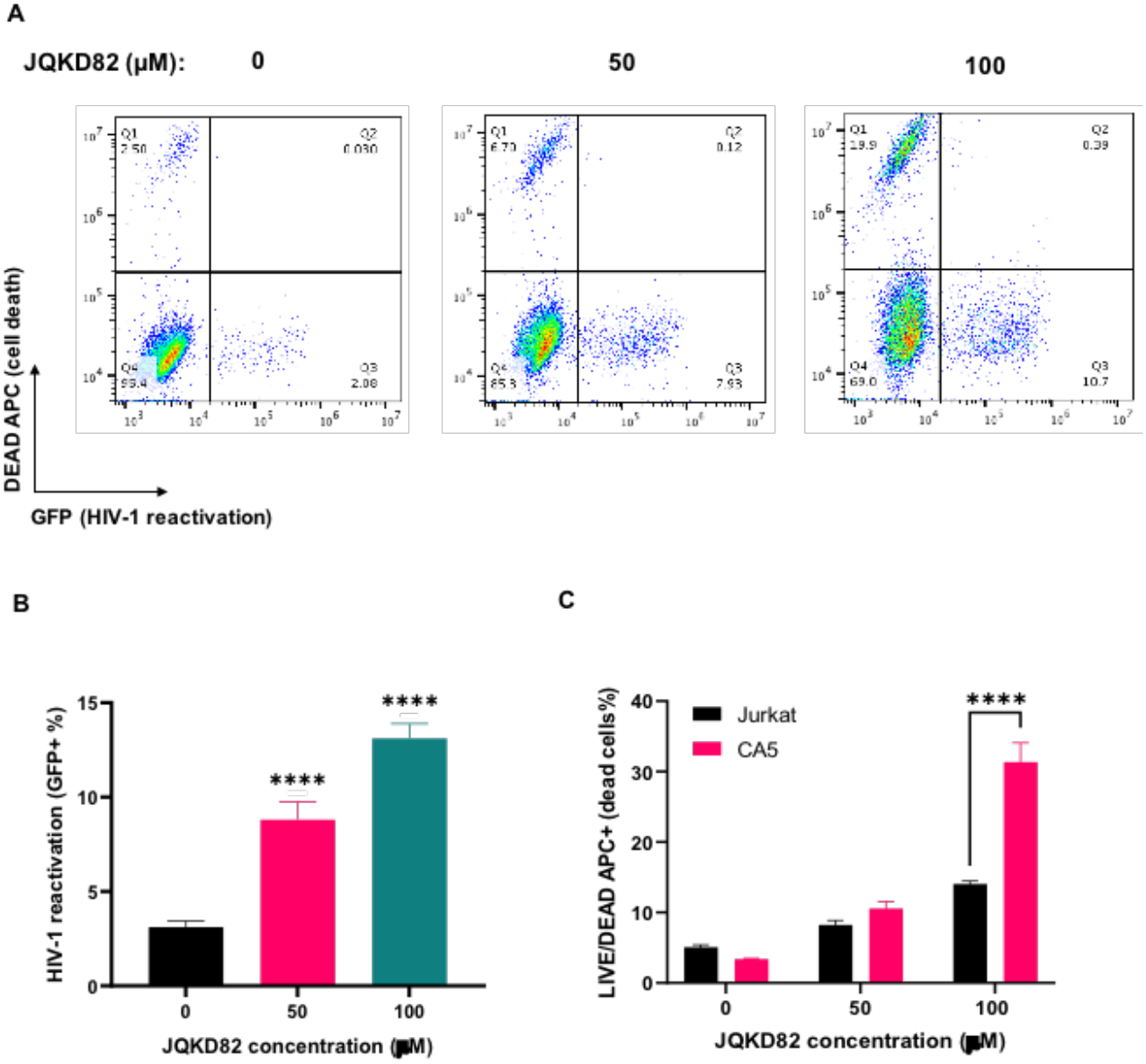
Short-time treatment of JQKD82 in CA5 cells for HIV-1 reactivation and cell killing. CA5 cells were treated with 0, 50, or 100 μM JQKD82 for 48 h. Cells were harvested for LIVE/DEAD staining and analyzed with FASC to identify the expression of HIV-1 LTR-driven GFP (**A, B**) and LIVE/DEAD-APC (**C**). Results were calculated from at least 3 independent experiments and presented as mean +/-standard error of the mean (SEM). (** p <0.01; **** p <0.0001 by one-way/two-way ANOVA and Tukey’s multiple comparison test compared to the untreated (**B**) or parental cell control group (**C**)).

**Fig S2.**
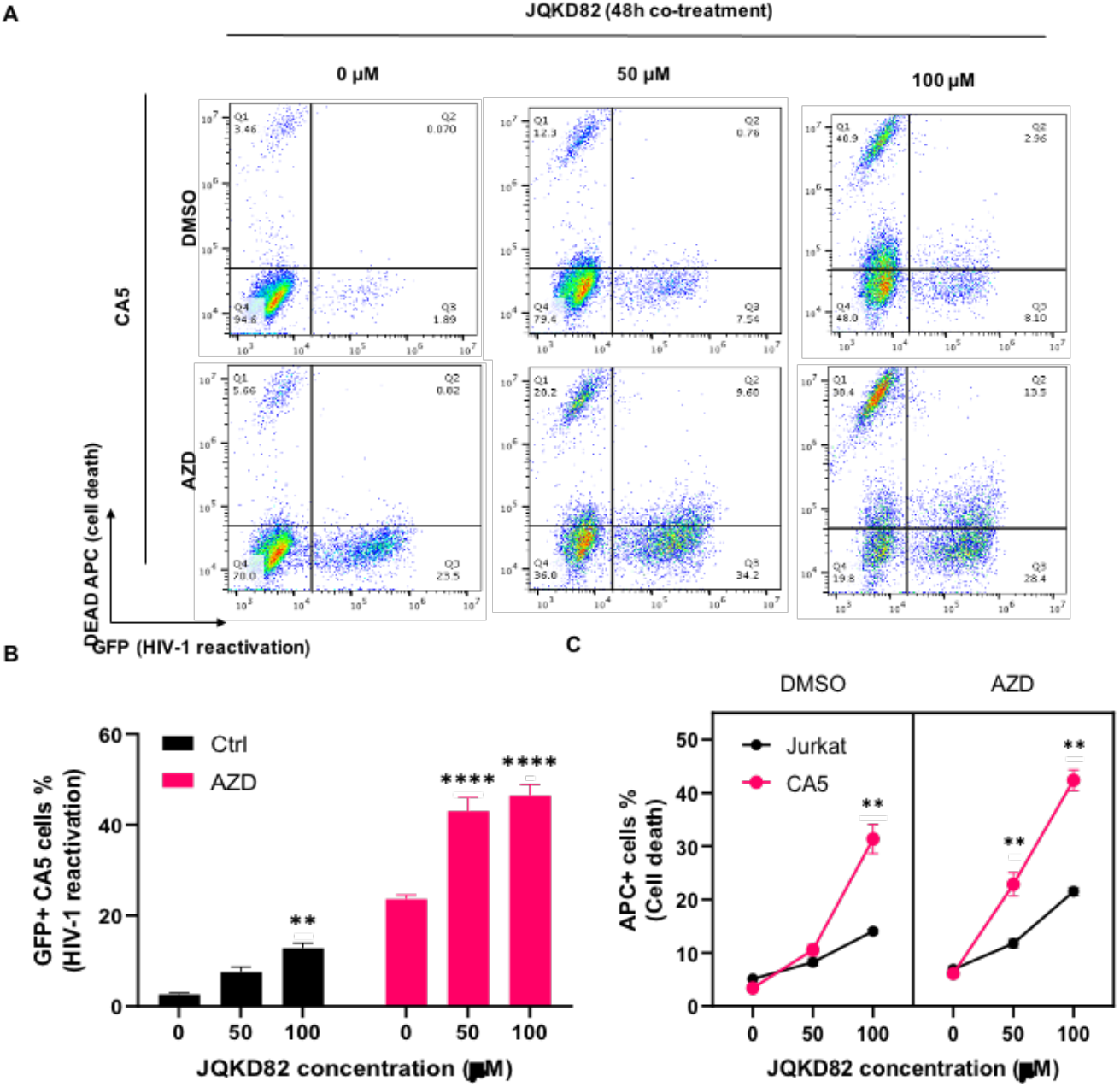
Short-time treatment of JQKD82/AZD5582 in CA5 cells for HIV-1 reactivation and cell killing. (**A**) CA5 cells were treated with 0, 50, or 100 μM with or without 0.2 μM AZD5582 for 48h. Cells were performed the LIVE/DEAD staining and analyzed by FASC. (**B**) Treated CA5 cells were analyzed by the GFP expression from the HIV-1 reactivation. Results were calculated from 3 independent experiments and were presented as mean +/-standard error of the mean (SEM). (** p <0.01; **** p <0.0001 by 2-way ANOVA and Tukey’s multiple comparison test compared to 0 μM JQKD82 treated group). (**C**) Treated CA5 cells were analyzed for the LIVE/DEAD APC expression. Results were calculated from 3 independent experiments and were presented as mean +/-standard deviation (SEM). (**p <0.01 by 3-way ANOVA and Tukey’s multiple comparison test compared to Jurkat cells under the same treatment.).

**Fig S3.**
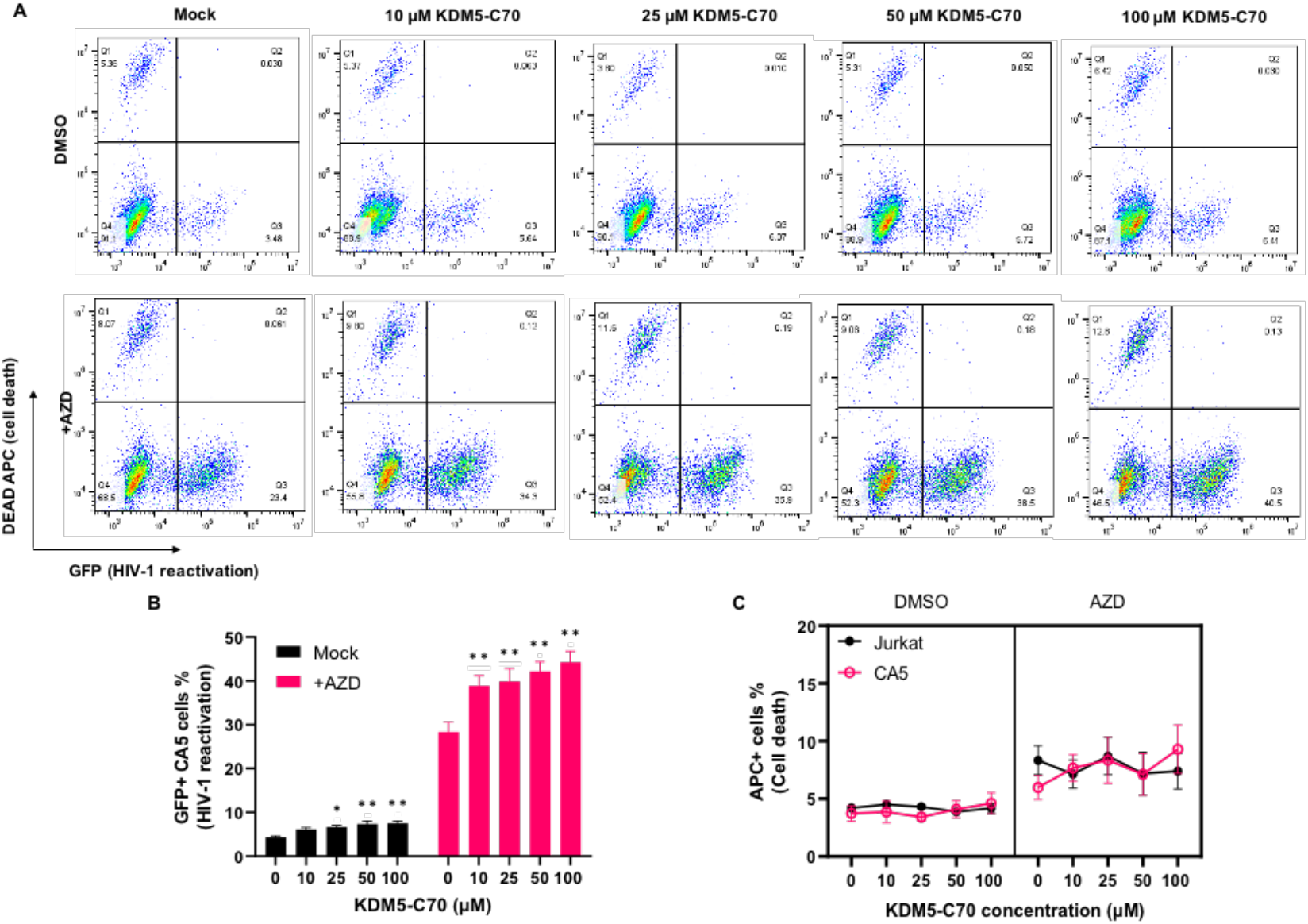
KDM5 inhibitor KDM5-C70 increased the HIV-1 reactivation in CA5 cells. (**A**) CA5 cells were treated with 0, 10, 25, 50, or 100 μM KDM5C-70 with or without 0.2 μM AZD5582 for 48h. Cells were performed the LIVE/DEAD staining and analyzed by FASC. (**B**) Treated CA5 cells were analyzed by the GFP expression from the HIV-1 reactivation. Results were calculated from 2 independent experiments and were presented as mean +/-standard error of the mean. (* p <0.05; ** p <0.01 by 2-way ANOVA and Tukey’s multiple comparison test compared to 0 μM KDM5-C70 treated group). (**C**) Treated CA5 and Jurkat parental cells were analyzed for the LIVE/DEAD APC expression. Results were calculated from 2 independent experiments and were presented as mean +/-standard error of the mean (SEM).

**Fig S4.**
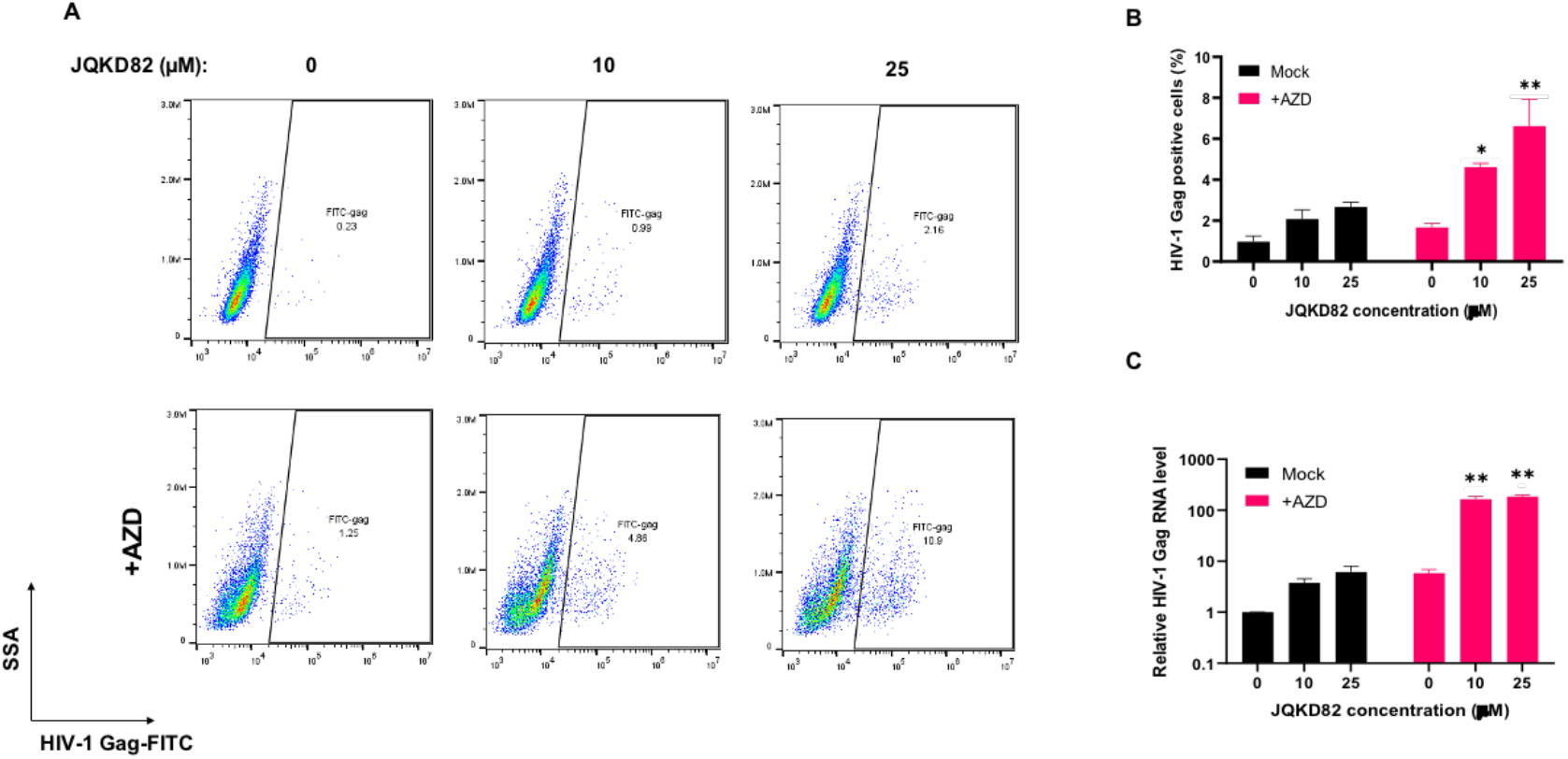
JQKD82/AZD5582 combinatory treatment increases the HIV-1 reactivation in U1/HIV monocyte cell line. (**A**) U1/HIV cells were treated with 0, 10, or 25 μM JQKD82 for 3 days and refreshed the treated medium with or without 0.2 μM AZD5582 for 48h. Cells were performed anti-HIV-1 Gag intracellular staining (**B**). Treated cells were harvested for RNA extraction and RT-qPCR to detect the HIV-1 Gag mRNA level (**C**). Results were calculated from at least 2 independent experiments and presented as mean +/-standard deviation (SD). (* p <0.05; ** p < 0.01; by two-way ANOVA and Tukey’s multiple comparison test compared to 0 μM JQKD82-treated control.)

